# Striatal and prefrontal D2R and SERT distributions contrastingly correlate with default-mode connectivity

**DOI:** 10.1101/2021.04.28.441769

**Authors:** Tudor M. Ionescu, Mario Amend, Rakibul Hafiz, Bharat B. Biswal, Andreas Maurer, Bernd J. Pichler, Hans F. Wehrl, Kristina Herfert

**Affiliations:** Werner Siemens Imaging Center, Department of Preclinical Imaging and Radiopharmacy, Eberhard Karls University Tuebingen, Germany; Department of Biomedical Engineering, New Jersey Institute of Technology, University Heights, Newark, New Jersey, USA

**Keywords:** Resting-State Functional Connectivity, Monoamines, D2 receptor, Serotonin Transporter, Simultaneous PET/fMRI

## Abstract

The molecular substrate of resting-state functional connectivity (rs-FC) remains poorly understood. We aimed to elucidate interactions of dopamine D2 receptor (D2R) and serotonin transporter (SERT) availabilities in main dopaminergic and serotonergic projection areas with the default-mode network (DMN) and two other resting-state networks (RSNs), the salience (SN) and sensorimotor networks (SMN). We performed simultaneous PET/fMRI scans in rats using [^11^C]raclopride and [^11^C]DASB to image D2R and SERT distributions, showing for the first time direct relationships between rs-FC and molecular properties of the rodent brain. We found negative associations between CPu D2R availability and all RSNs investigated. Strikingly, medial prefrontal SERT correlated both positively with anterior DMN rs-FC and negatively with rs-FC between the other networks, underlining serotonin’s intricate role in this region. By further elucidating the link between molecular brain properties and its network-level function, our data support future diagnostic and therapeutic strategies.

**Teaser:** Simultaneous PET/fMRI indicates direct associations between monoaminergic neurotransmission and brain functional networks.

## Introduction

Resting-state functional connectivity (rs-FC) derived from functional magnetic resonance imaging (fMRI) studies has emerged as a promising biomarker to assess brain function and dysfunction over the last two decades [1]. While rs-FC has already been proven to be a valuable tool for the basic understanding of brain pathology [2], there are still many unresolved aspects regarding its emergence and modulation. One important, still not elucidated question is the modulatory role of neurotransmitters and receptors on rs-FC. In most studies, pharmacological MRI (ph-MRI) has been used to investigate the impact of neurotransmitters on rs-FC [3, 4]. However, its output reflects the active pharmacological manipulation of the entire brain rather than the effects of intrinsic regional dependencies between neurotransmitter signaling and rs-FC. To this extent, PET/fMRI studies are a powerful approach to investigate brain network modulation by different neurotransmitter systems [5-9]. However, only a few studies have employed simultaneous PET/fMRI, a prerequisite for an accurate temporal and spatial cross-correlation [5, 7].

Here, we present the first simultaneous PET/fMRI approach in rats to delineate the relations between regional D2 receptor (D2R) and serotonin transporter (SERT) availabilities studied with PET and RSNs studied with fMRI. Recently, Conio et al. revealed opposing roles of dopamine (DA) and serotonin (5-HT) on three human RSNs: the default-mode network (DMN), postulated to be involved in functions such as self-reference, memory formation and imagination [10], the sensorimotor network (SMN), regulating sensory processing [11] and the salience network (SN), playing important roles in salience attribution and reward and being assumed to mediate the interplay between the DMN and task-positive networks such as the SMN [12, 13].

In the study by Conio et al., the authors investigated the effects of monoaminergic synthesis in the raphé nuclei and substantia nigra on the mentioned RSNs [14]. Here, we selected the same three RSNs for investigation in our study, focusing on the DMN, due to its prominent role in different pathologies [2, 11, 15]. However, in contrast to the paper by Conio et al., we aimed to elucidate the correlations of D2R and SERT distributions in the caudate putamen (CPu) and medial prefrontal cortex (mPFC), two of the most prominent dopaminergic and serotonergic projection areas. Our preclinical data acquired using uniform cohorts regarding age, strain, gender, nutrition, and living conditions are likely to reflect correlations driven by intrinsic differences between individual subjects. We chose to focus on DA and 5-HT due to their modulatory role in several important brain functions, such as motor control, motivation, mood, and emotion and thus their involvement in different neurodegenerative and psychiatric diseases such as Parkinson’s disease (PD) and major depressive disorder (MDD) [14, 16, 17]. Insight into the intrinsic correlations of D2R and SERT with rs-FC may improve therapy and drug development for such pathologies.

## Results

The results indicate inter-subject correlations between rs-FC and [^11^C]raclopride and [^11^C]DASB BP_nd-norm_ values in the CPu and mPFC. The significances depicted were calculated at p < 0.05 (both uncorrected and FDR-corrected using Benjamini-Hochberg, please refer to corresponding figure legends). For more detailed information, correlations with significances of p < 0.01 and p < 0.001 are indicated in the *Supplementary Information* for all presented matrices.

### D2 receptor availability in CPu correlates with widespread reduced rs-FC

Due to the importance of dopamine in the CPu in modulating brain function, we aimed to elucidate the intrinsic correlation of D2R availability in this region to the rs-FC of the DMN, SN and SMN.

Increased D2R availability was associated with reduced rs-FC between all three analyzed networks, as depicted in Figure 2. On edge level (Figure 2A), most significant correlations involved the mPFC (23 decreased edges) and PaC (35 decreased edges) in the DMN, Cg (21 decreased edges) and Ins (29 decreased edges) in the SN and the CPu (27 decreased edges), M1 (36 decreased edges) and S1 (36 decreased edges) in the SMN.

**Figure 1:**
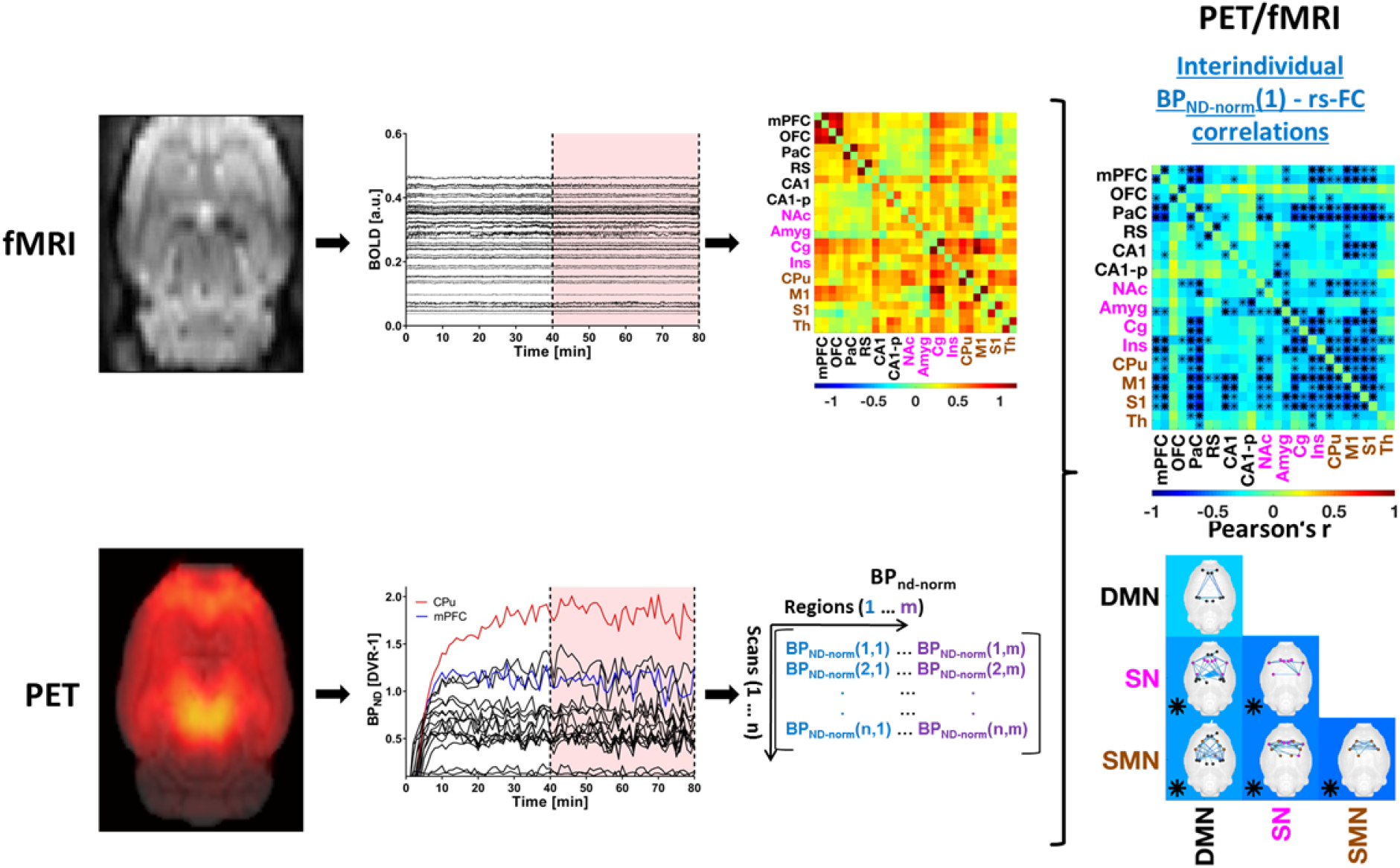
PET/fMRI data analysis following preprocessing. fMRI: regional BOLD time-courses were extracted from each scan and subject and rs-FC matrices were computed. **PET:** regional DVR-1 values were extracted from 40 to 80 min after tracer injection for each scan and subsequently normalized to whole brain values. **PET-fMRI:** For every region the correlations of its subject-wise BP_ND-norm_ values and every subject-wise rs-FC edge were calculated, resulting in inter-individual correlation matrices per region between respective PET tracer binding and rs-FC.

**Figure 2:**
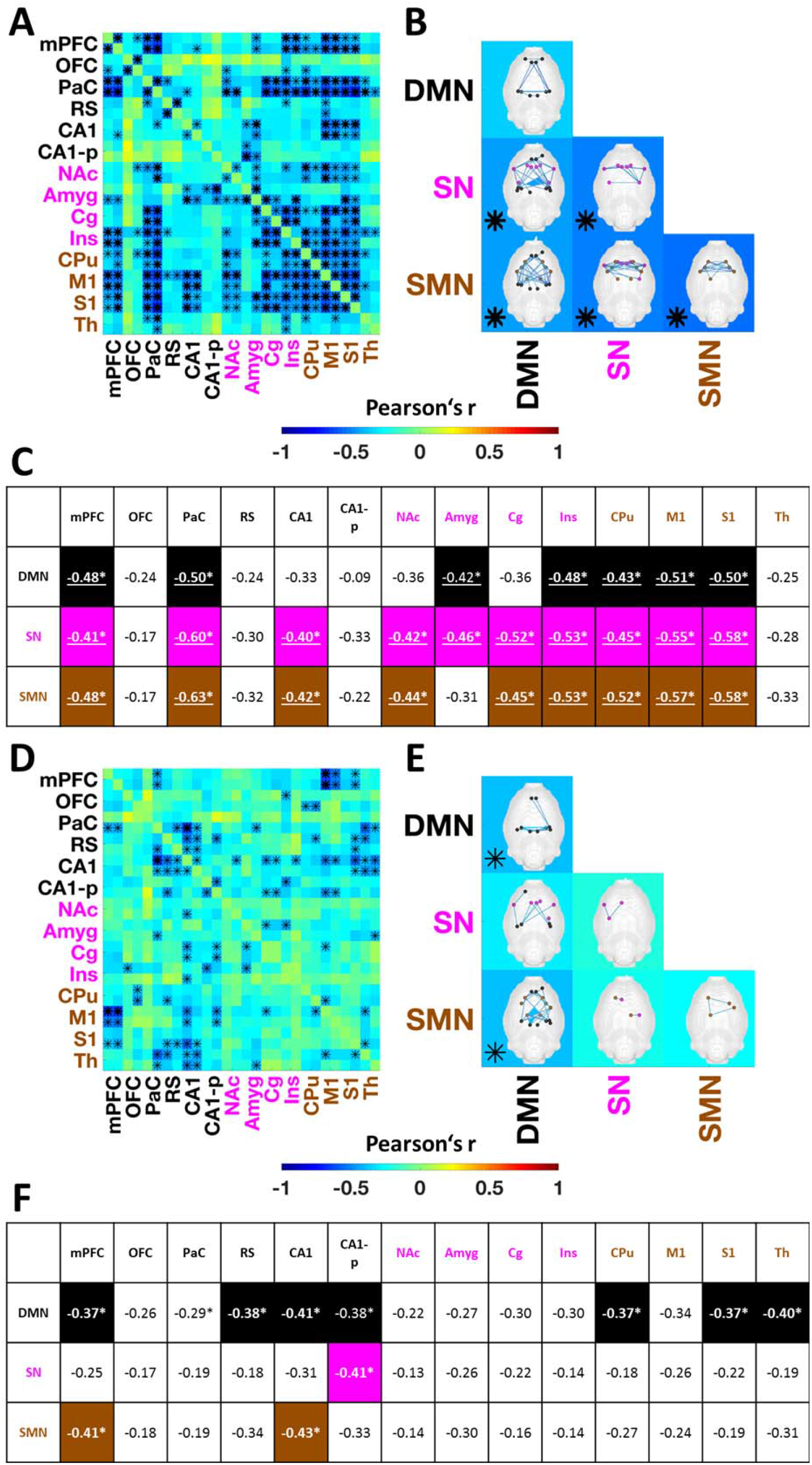
Correlations between D2R availability in the CPu and mPFC and rs-FC. **(A)** The correlation matrix indicates all correlations between CPu [^11^C]raclopride BP_ND-norm_ and pairwise rs-FC values between regions comprising the DMN, SN and SMN. DMN regions are depicted in black, SN regions in magenta and SMN regions in brown color. * indicates statistically significant correlations (p < 0.05, bold asterisks indicate correlations surviving FDR correction). **(B)** Matrix indicating correlations between CPu [^11^C]raclopride BP_ND-norm_ and within-network or between-network connectivity strengths for the DMN, SN and SMN. * indicates statistically significant correlations (p < 0.05, bold asterisks indicate correlations surviving FDR correction). The brain maps depict the within-network or between-network edges significantly correlating with CPu BP_ND-norm_ values. **(C)** Table indicating the correlations (Pearson’s r) of CPu [^11^C]raclopride BP_ND-norm_ and the rs-FC strengths of each analyzed region (averaged between left and right hemisphere) to the DMN, SN and SMN. Significant correlations are highlighted by the underlying color (DMN – black, SN – magenta, SMN -brown) and asterisks. Correlations surviving FDR correction are underlined. **(D)** The correlation matrix indicates all correlations between mPFC [^11^C]raclopride BP_ND-norm_ values and pairwise rs-FC values between regions comprising the DMN, SN and SMN. DMN regions are depicted in black, SN regions in magenta and SMN regions in brown color. * indicates statistically significant correlations (p < 0.05, bold asterisks indicate correlations surviving FDR correction). **(E)** Matrix indicating correlations between mPFC [^11^C]raclopride BP_ND-norm_ and within-network or between-network connectivity strengths for the DMN, SN and SMN. * indicates statistically significant correlations (p < 0.05, uncorrected). The brain maps depict the within-network or between-network edges significantly correlating with mPFC BP_ND-norm_ values. **(F)** Table indicating the correlations of mPFC [^11^C]raclopride BP_ND-norm_ and the rs-FC strengths of each analyzed region (averaged between left and right hemisphere) to the DMN, SN and SMN. Significant correlations are highlighted by the underlying color (DMN – black, SN – magenta, SMN -brown) and asterisks (p < 0.05, uncorrected). Abbreviations: D2R = D2 receptor, BP_ND-norm_ = normalized binding potential, rs-FC = resting-state functional connectivity, CPu = caudate putamen, mPFC = medial prefrontal cortex, DMN = default-mode network, SN = salience network, SMN = sensorimotor network. For a list of abbreviations of all regions please refer to *Supplementary Table 1*.

SN and SMN within-network strengths (Figure 2B) were anti-correlated to the D2R availability in the CPu (r = -0.52 (p < 0.05, FDR-corrected) for SN and r = -0.53 (p < 0.05, FDR-corrected) for SMN). Similarly, the between-network connectivity was decreased for each pair of networks (r = -0.45 (p < 0.05, FDR-corrected) between DMN and SN, r = -0.46 (p < 0.05, FDR-corrected) between DMN and SMN; r = -0.51 (p < 0.05, FDR-corrected) between SN and SMN).

The correlations calculated for within-network and between-network node strengths and CPu D2R availabilities also included significant values for all three networks (Figure 2C). The connectivity strengths of six regions, including mPFC, PaC, Ins, CPu, M1 and S1 to all three networks were significantly anti-correlated to D2R binding the striatum (p < 0.05, FDR-corrected). The strongest negative correlations were observed for the SMN, ranging up to r = -0.63 for the rs-FC of PaC, a total of nine regions being negatively correlated with the SMN at a significance threshold of p < 0.05 including FDR correction (r = -0.48 for mPFC, r= -0.42 for CA1, r = -0.44 for NAc, r = -0.45 for Cg, r = -0.53 for Ins, r = -0.52 for CPu, r = -0.57 for M1, r = -0.58 for S1). The node strengths of the same regions to the SN, with the exception of the Amyg, also correlated with CPu [11C]raclopride binding at p < 0.05 with FDR correction, the correlation coefficients ranging up to r = -0.60 for PaC and r = -0.58 for S1. The node strengths of seven regions to the DMN correlated significantly (p < 0.05, FDR correction) with CPu D2R density (ranging from r = -0.42 for Amyg to r = -0.51 for M1).

### Medial prefrontal D2R correlates negatively with rs-FC of regions involved in cognitive control

We investigated the relationships between medial prefrontal D2R availability and rs-FC due to the reported role of dopamine in cognitive control mediated by this brain region.

Compared to the widespread relationships between D2R binding in the CPu and rs-FC (Figure 2A-C), the correlations observed between medial prefrontal [^11^C]raclopride BP_ND-norm_ values and the rs-FC of the three analyzed RSNs were sparse and to a large extent involved the DMN (Figure 2D-F). Specifically, 45 of the 52 edges significantly anti-correlated to D2R availability involved at least one region belonging to the DMN (Figure 2D). The strongest correlations occurred for the edges between mPFC and MC (up to r = -0.53, p < 0.05, FDR-corrected).

On network level (Figure 2E), the medial prefrontal D2R availability was significantly anti-correlated with the within-network DMN rs-FC strength (r = -0.40, p < 0.05), as well as the rs-FC between DMN and SMN (r = -0.40, p < 0.05). Connectivity strengths within and between SN and SMN did not correlate significantly with medial prefrontal D2R binding.

The within-network strengths of three posterior DMN regions were associated with increased D2R binding in the mPFC (Figure 2F), including RS (r = -0.38, p < 0.05), CA1 (r = -0.41, p < 0.05) and CA1-p (r = -0.38, p < 0.05). In addition, decreased connectivity to the DMN was detected in the CPu (r = -0.37, p < 0.05), belonging to as well as in the S1 (r = -0.37, p < 0.05) and Th (r = -0.40, p < 0.05), all regions being part of the SMN. In contrast, only the rs-FC of CA1-p (r = -0.41, p < 0.05) to the SN, as well as the rs-FC of mPFC (r = -0.41, p < 0.05) and CA1 (r = -0.43, p < 0.05) correlated negatively with increased medial prefrontal D2R availability.

### SERT availability in the CPu negatively correlates with SN connectivity

Correlations of SERT availability in the CPu with rs-FC were subtle and mainly restricted to SN connectivity. Specifically, negative correlations were found between SERT availability and 7 out of 28 edges in the SN. Five edges of the NAc and Ins respectively to other SN regions were significantly decreased (p ≤ 0.05, Figure 3A). Additionally, NAc and Ins edges to regions outside the SN including CPu, S1 and and RS correlated negatively with SERT availability in the CPu. Finally, 3 out of 4 edges between the CPu itself and S1 correlated negatively with SERT availability. The sparse edge-wise correlations did not however translate to correlations with either within or between-network rs-FC (Figure 3B). Nonetheless, the integration of NAc and Ins within the SN correlated negatively with SERT availability in the CPu (p < 0.05, Figure 3C). Interestingly, multiple edges involving DMN rs-FC correlated positively, though not significantly with SERT density in the striatum, in contrast to the negative correlations observed between striatal SERT availability and salience rs-FC and to a lesser extent SMN rs-FC.

**Figure 3:**
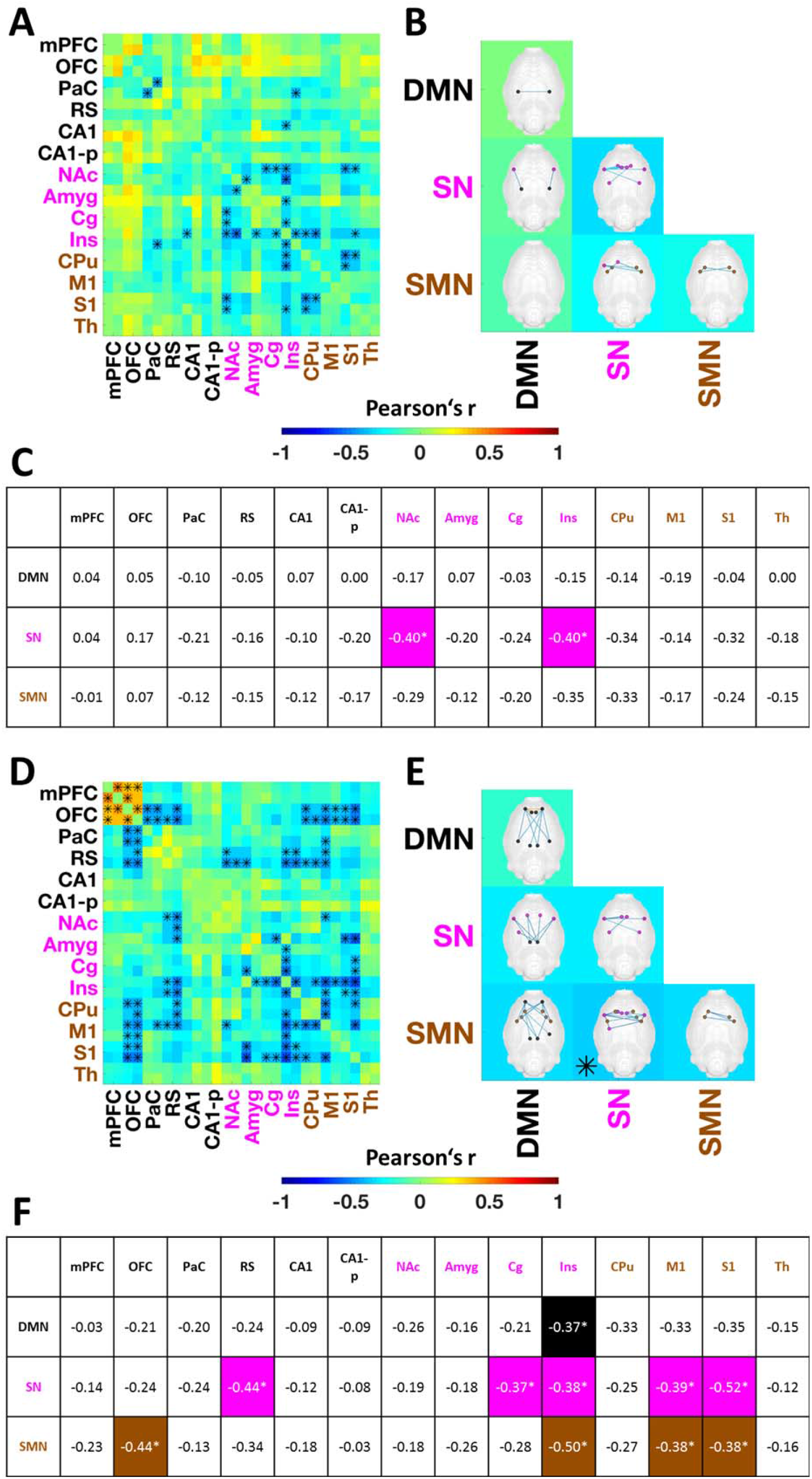
Correlations between SERT availabilities in the CPu and mPFC and rs-FC. **(A)** The correlation matrix indicates all correlations between CPu [^11^C]DASB BP_ND-norm_ values and pairwise rs-FC values between regions comprising the DMN, SN and SMN. DMN regions are depicted in black, SN regions in magenta and SMN regions in brown color. * indicates statistically significant correlations (p < 0.05, uncorrected). **(B)** Matrix indicating correlations between CPu [^11^C]DASB BP_ND-norm_ and within-network or between-network connectivity strengths for the DMN, SN and SMN. * indicates statistically significant correlations (p < 0.05, uncorrected). The brain maps depict the within-network or between-network edges significantly correlating with CPu [^11^C]DASB BP_ND-norm_ values. **(C)** Table indicating the correlations of CPu [^11^C]DASB BP_ND-norm_ and the rs-FC strengths of each analyzed region (averaged between left and right hemisphere) to the DMN, SN and SMN. Significant correlations are highlighted by the underlying color (DMN – black, SN – magenta, SMN -brown) and asterisks (p < 0.05). **(D)** The correlation matrix indicates all correlations between mPFC [^11^C]DASB BP_ND-norm_ values and pairwise rs-FC values between regions comprising the DMN, SN and SMN. DMN regions are depicted in black, SN regions in magenta and SMN regions in brown color. * indicates statistically significant correlations (p < 0.05, uncorrected). **(E)** Matrix indicating correlations between mPFC [^11^C]DASB BP_ND-norm_ and within-network or between-network connectivity strengths for the DMN, SN and SMN. * indicates statistically significant correlations (p < 0.05, uncorrected). The brain maps depict the within-network or between-network edges significantly correlating with mPFC [^11^C]DASB BP_ND-norm_ values. **(F)** Table indicating the correlations of mPFC [^11^C]DASB BP_ND-norm_ and the rs-FC strengths of each analyzed region (averaged between left and right hemisphere) to the DMN, SN and SMN. Significant correlations are highlighted by the underlying color (DMN – black, SN – magenta, SMN -brown) and asterisks (p < 0.05, uncorrected). Abbreviations: SERT = serotonin transporter, BP_ND-norm_ = normalized binding potential, rs-FC = resting-state functional connectivity, Cpu = caudate putamen, mPFC = medial prefrontal cortex, DMN = default-mode network, SN = salience network, SMN = sensorimotor network. For a list of abbreviations of all regions please refer to *Supplementary Table 1*.

### Prefrontal SERT specifically increases anterior DMN rs-FC

Expressing the highest [^11^C]DASB binding of the entire cortex, we evaluated the mPFC to elucidate whether its SERT availability correlates with the rs-FC of the DMN, SN and SMN (Figure 3D-F).

The correlations of the investigated edges with medial prefrontal SERT availability were heterogeneous (Figure 3D). Specifically, the short-range rs-FC within the anterior DMN comprised of mPFC and OFC correlated positively with [^11^C]DASB binding, reaching up to r = 0.45 (p < 0.05) between left and right mPFC. In contrast, long-range rs-FC of the OFC was anti-correlated to mPFC SERT within the DMN to PaC and RS (r = -0.49, p < 0.01 between right OFC and right RS) as well as to the SMN regions CPu (r = -0.43, p < 0.05 between left OFC and CPu), M1 and S1 (r = -0.5, p < 0.01 between right OFC and right S1). Further negative correlations were found for edges between RS and regions belonging to the SN, such as NAc (r = -0.45, p < 0.05 for right RS – left NAc), Amyg (r = -0.41, p < 0.05 for right RS – left Amyg), or Ins (r = -0.5, p < 0.01 for right RS – right Ins). Finally, several edges involving both SN and SMN nodes were significantly reduced at higher medial prefrontal SERT availabilities. Most prominently, 16 edges of the Ins to regions including RS, Amyg, Cg, CPu, M1 and S1 were decreased. Additionally, mPFC [^11^C]DASB binding was negatively correlated with 14 S1 edges to regions such as OFC, Amyg, Cg, Ins, CPu and M1.

On a network level the rs-FC between SN and SMN was reduced significantly (r = -0.36, p < 0.05) by increasing SERT availability in the mPFC (Figure 3E). The rs-FC between and within the other RSNs was not significantly associated to SERT levels in this region. Finally, Figure 3C indicates significant correlations between region-wise rs-FC to the three RSNs and medial prefrontal SERT availability. The Ins was the only region with significantly anti-correlated rs-FC to the DMN (r = -0.37, p < 0.05) and the only node with significantly decreased rs-FC to all three RSNs (r = 0.38, p < 0.05 to the SN and r = 0.5, p < 0.01 to the SMN) associated with increased medial prefrontal SERT availability. Five regions, including RS, Cg, M1 and S1 in addition to Ins had significantly lower rs-FC strength to the SN, while OFC (r = -0.44, p < 0.05), M1 and S1 (both r = -0.38, p < 0.05) were significantly less connected to the SMN at increasing [^11^C]DASB bindings in the mPFC.

We further elucidated the way prefrontal SERT binding shifts the rs-FC balance towards the anterior DMN, by calculating the correlation of medial prefrontal [^11^C]DASB binding with the difference of the average rs-FC within the anterior DMN and the average rs-FC between the anterior DMN and the posterior DMN, SN and SMN (please refer to *Supplementary Information*). This analysis yielded a highly significant correlation value of r = 0.63 (p = 0.0003), emphasizing the shift in processing balance by the corroborated positive correlation of the anterior DMN within-network rs-FC with the negative correlation of the anterior DMN between-network rs-FC with medial prefrontal SERT binding.

## Discussion

Elucidating the link between molecular variations of brain receptors and transporters and macroscale metrics such as rs-FC will primarily enhance our understanding about drug mechanisms of action and several brain pathologies. Here we show that D2R and SERT availabilities correlate with rs-FC in a regionally specific manner in the healthy rat brain (see Figure 4 for a summary of the findings).

**Figure 4:**
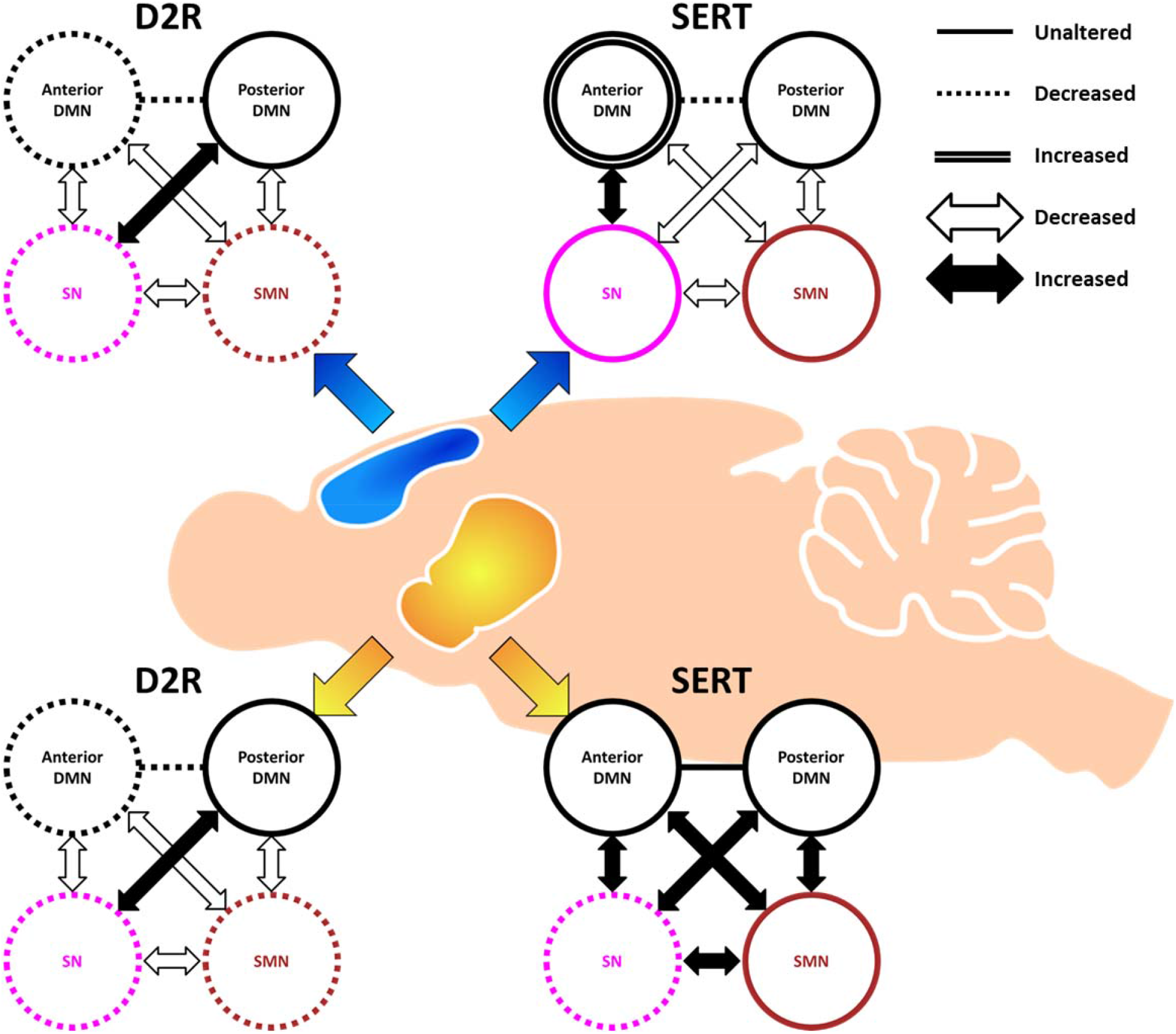
Summary of the findings. In the mPFC (blue) D2Rs are negatively correlated with posterior DMN rs-FC and the rs-FC between DMN and SMN. SERT availability in the mPFC is negatively correlated with rs-FC between anterior and posterior DMN, yet positively correlated with anterior DMN rs-FC. The rs-FC between SN and SMN is also anti-correlated with medial prefrontal SERT density. In the CPu (yellow) increasing D2R was associated with decreased rs-FC within and between all networks except for the largely unaffected posterior DMN, while SERT availability correlated negatively with SN rs-FC and did not correlate to rs-FC in other networks. Interrupted lines indicate negatively correlated within-network rs-FC, empty arrows indicate negatively correlated between-network rs-FC. Single continuous lines and full arrows indicate no correlation of within and between-network rs-FC. Double lines indicate positive correlation of within-network rs-FC. Abbreviations: D2R = D2 receptor, SERT = serotonin transporter, DMN = default-mode network, SN = salience network, SMN = sensorimotor network.

### D2 receptor density in the CPu is negatively correlated with rs-FC across all RSNs

DA and the mesolimbic system in particular, expressing the highest density of dopaminergic receptors, are of paramount importance for numerous neurological disorders for which rs-FC can serve as a biomarker [18]. The D2Rs investigated in the present study using [^11^C]raclopride drive the indirect striatal pathway, a circuit involved in the inhibition of motor activity via the ventrolateral nucleus of the thalamus postulated to be heavily involved in PD [19, 20]. PET studies using [^11^C]raclopride have been shown to provide an indirect measure of synaptic dopamine availability with a decrease in [^11^C]raclopride binding reflecting an increase in synaptic dopamine concentrations due to a higher occupancy at the receptor and vice versa [21]. Thus, a higher D2R availability reflects a lower synaptic dopamine content, which may impact rs-FC in downstream pathways. The largest decrease associated with higher D2R availability in the caudate putamen observed in our study occurred in the SMN, in line with the increased activity of the indirect pathway mediated by higher D2R densities. However, our data also suggests reduction in DMN and SN rs-FC. Regions comprising these networks may be mediated along different pathways by striatal D2Rs, as shown previously [22]. Specifically, striatal D2R overexpression in mice has been shown to modulate ventral tegmental activity [23]. One of the effects of this modulation is the impairment of functional connectivity between VTA and mPFC, resulting in abnormal prefrontal processing and affecting working memory [24]. Intriguingly, the connectivity between the medial prefrontal cortex and other areas correlated strongly to D2R availability in the caudate putamen in our study, in line with the reports discussed above.

To the best of our knowledge, the only simultaneous PET/fMRI study up to date which explored the correlation of rs-FC with receptor variability employed [^11^C]NNC112, a dopamine D1 receptor PET ligand in the human brain [5]. The primary finding of this study was a correlation between D1R cortical density and the functional connectivity between DMN and the frontoparietal network during a working memory task. Intriguingly, the authors also found a significant negative correlation between striatal D1R availability and rs-FC between left and right mPFC. While our data reveal large-scale decoupling of the mPFC from several other regions at increased striatal D2R densities, the rs-FC between left and right mPFC did not correlate with D2R in the striatum. These findings indicate that future studies assessing both D1R and D2R availability are required to accurately delineate their combined interaction with rs-FC.

Another study, applying PET and fMRI sequentially rather than simultaneously in humans, focused on the effects of both DA synthesis using [^18^F]DOPA and release capacity using [^11^C]-(+)-PHNO on rs-FC in placebo and dexamphetamine challenge studies [6]. While [^18^F]DOPA PET as a marker of presynaptic aromatic amino acid decarboxylase (AADC) activity is not linked to a particular receptor subtype, [^11^C]-(+)-PHNO binds specifically to D2/3 receptors, similarly to [^11^C]raclopride in our study. The data showed that DA release capacity to D2/3 receptors in the CPu was negatively correlated to the salience network connectivity, which is in line with our findings, indicating the translatability of this study design among species. Interestingly, in the above-mentioned study DA synthesis capacity correlated with increased salience rs-FC. The authors corroborated their findings to the hypothesis that DA synthesis reflects a general dopaminergic tone that would be necessary for the attribution of salience, while DA release would indicate spontaneous stimulus-independent firing mediating aberrant attributions of salience. Our data thus support the mentioned hypothesis. Additionally, the authors also discussed the possibility of the preferential binding of the agonist [^11^C]-(+)-PHNO tracer they used to high-affinity D2R [25] having an effect on their readout. To this extent they proposed the use of an antagonist tracer for further elucidation of this aspect. The similarity of our readout using [^11^C]raclopride, a D2 antagonist tracer, complements the findings using [^11^C]-(+)-PHNO indicating that the proportion of high-affinity D2R does not have a major impact on the readout in this case.

### Medial prefrontal D2R availability is associated with reduced DMN connectivity

Several reports have suggested an essential role of DA in the prefrontal cortex, an associative cortical area involved in the top-down control of several cognitive mechanisms [26]. The balance between D1 and D2 receptors available in this region has been linked to normal brain function and is postulated to play a critical role in psychiatric diseases such as schizophrenia [27]. It is assumed that D1 and D2 receptors play complementary roles in functions such as associative learning, with D2R promoting cognitive flexibility. Thus, increased prefrontal D2R availability destabilizes network states promoting flexible behavior [28-30]. This hypothesis is supported by our findings, indicating a decrease in rs-FC, especially in the DMN, a network postulated to be heavily involved in cognitive control.

### Striatal SERT subtly impacts salience network circuitry

While serotonergic innervation in the striatum by the raphé nuclei has been demonstrated [31], the role of serotonin in the striatum remains largely elusive. Early studies have indicated a dose-dependent interaction between exogenous serotonin and dopaminergic activity [32]. However, when compared to the concentrations of at least 100 nM assessed as having an effect on dopaminergic activity [33], the endogenous levels of serotonin determined in the striatum at resting state (0.5 – 2 nM) appear insufficient to directly impact dopaminergic activity in this region [32], which may explain the relatively sparse correlations observed here between SERT density and rs-FC. Nonetheless, other studies have suggested that the effect of serotonin in the striatum may be mediated by additional factors, such as a state dependence of dopaminergic neurotransmission [32] or by interactions with other neurotransmitter systems altogether [34].

The hypothesis indicating a subtle, yet physiologically significant role of serotonin in the striatum is supported by macroscale findings, including those generated by rs-FC studies. Specifically, decreased FC between the raphé nuclei and the striatum has been associated with decreased connectivity of the salience network with subcortical regions in schizophrenia [35]. The significant effects observed in the present study mirror the findings by Han et al. [35], being largely confined to salience connectivity. Potential pathological roles of such correlations include the postulated aberrant salience attribution in schizophrenia, as well as the reported deficient motivation [35], a hypothesis supported by the significantly decreased rs-FC strength of the nucleus accumbens within the salience network found in our study. Intriguingly, similar findings of decreased salience rs-FC have been associated with increased D2 availability in the striatum [36], which is also confirmed by the findings in the present study. Therefore, our data indicate that striatal D2R and SERT densities have similar effects on the salience network in particular, both being anti-correlated with its rs-FC. However, two aspects must be underlined. First, our data indicate that striatal D2R correlations with rs-FC are stronger than those of SERT and also involve the two other investigated networks. As a side note, although not achieving significance, the positive correlations observed between striatal SERT and anterior DMN rs-FC in the present study antagonize its negative correlations with salience rs-FC and suggest that the role of serotonin in this region may be network-dependent. Secondly, potential direct interactions between SERT and D2R in the striatum have not been elucidated in the present study. Due to the similar effects of D2R and SERT underlined above, future studies assessing both parameters along with rs-FC in the same cohort are of interest to elucidate potential three-way interactions.

### Medial prefrontal SERT density has opposing effects on anterior and posterior default-mode connectivity

Our data indicate that SERT availability in the mPFC has a heterogeneous impact on rs-FC, localized positive correlations in the anterior DMN being corroborated with more widespread negative correlations in the SN and SMN.

The associations observed in the present study could provide important insights for MDD and related disorders, which are associated with disrupted serotonergic neurotransmission. In MDD, cortical 5-HT levels are believed to be decreased, one of the possible causes being an increased expression of SERT [37-39]. Due to the rostro-caudal gradient in serotonergic innervation it is widely postulated that 5-HT in the prefrontal cortex may play an essential role in rs-FC modulation and thereby in numerous psychiatric diseases including MDD [26, 40]. In MDD, most rs-FC research has focused on the DMN, a network associated with a state of enhanced rumination. Taken together, past studies indicate increased rs-FC in frontal areas corroborated with decreases in the posterior default-mode hubs [15]. In line with these findings, we demonstrate an increased local rs-FC within the prefrontal cortex associated at elevated medial prefrontal SERT availability and concurrent with decreased rs-FC in the posterior DMN, as well as in the SMN and SN.

Further evidence indicating the prominent role of the prefrontal cortex in depression is provided by acute tryptophan depletion (ATD) [41, 42] and selective serotonin reuptake inhibitors (SSRI) studies. Briefly, both ATD and single-dose SSRI administration have been shown to increase local prefrontal rs-FC by decreasing serotonergic levels [43, 44]. In contrast, chronic SSRI medication reduced pathologically altered rs-FC of the medial prefrontal cortex in MDD [45, 46]. Our data indicating an increased local prefrontal rs-FC and decreased rs-FC in posterior DMN, as well as in the SN and SMN at higher prefrontal SERT levels are in line with previous studies. A localized prefrontal rs-FC increase with its concurrent dissociation from most other areas of the brain may indicate an enhanced state of rumination corroborated with a loss of top-down control and regulation [46-50]. Another interesting aspect of our data is the decoupling of the insular cortex from all RSNs and most strongly from the task-positive SMN at increased prefrontal SERT densities. Being the main hub of the salience network, this finding may suggest that disturbed regulation of DMN-SMN balance, one of the main functions of the SN [14, 51], is at least in part associated with prefrontal serotonergic function.

Using sequential [^11^C]WAY-100635 PET/fMRI Hahn et al. investigated the modulations of the DMN by regional 5-HT_1A_ receptor availabilities [8]. The authors found reduced DMN rs-FC at increased 5-HT_1A_ receptor binding in the dorsal medial prefrontal cortex, in line with our results using [^11^C]DASB. However, when comparing the study by Hahn et al. with the present study, it should be kept in mind that Hahn et al. investigated correlations between rs-FC and a single 5-HT receptor subtype. The effects of 5-HT are mediated via at least 14 5-HT receptor subtypes, investigating a single 5-HT receptor subtype provides only one possible modulation of the rs-FC elicited by 5-HT [26], while imaging SERT availability as done in our study may represent a more general reflection of regional serotonergic tone. The two approaches should be seen as complementary and future studies investigating the influence of both SERT and different 5-HT receptors on rs-FC will further enhance our understanding of the serotonergic system.

### Serotonergic and dopaminergic correlations with rs-FC

Our data shows that individual variations of regional D2R and SERT availabilities at rest correlate with different aspects of the analyzed RSNs. The recent study by Conio et al. indicated specific, mainly opposite roles of the two neurotransmitters in the modulation of DMN, SN and SMN [14]. However, the authors mainly focused on the correlation between rs-FC and neurotransmitter synthesis at the raphé nuclei and substantia nigra. Our study complements the proposed model by showing that the availabilities of receptors in projection areas also play essential roles in the way the respective neurotransmitters modulate rs-FC. Specifically, we found that the rs-FC of main functional hubs, well-connected regions known to both receive projections and send afferents to widely distributed brain areas correlate strongest with their respective D2R or SERT availabilities. The CPu is a main hub of the basal ganglia [52], known to modulate motor functions via the cortico-striato-thalamic loop [11], but also salience and prefrontal function via the VTA [24]. The mPFC is the main hub of working memory and attention, integrating inputs from multiple sensory modalities [47]. Moreover, monoaminergic function in these regions is at the center of various diseases. Dopamine in the caudate putamen plays an essential role in PD [20], while striatal SERT has been reported to play an important role in schizophrenia. Medial prefrontal serotonergic dysfunction is related to MDD [53], and dopaminergic imbalance in the mPFC is postulated to drive schizophrenia [54]. Our data show the importance of monoamines in these hubs not only for their own function but for the modulation of the most important RSNs of the brain. Additionally, the complementing associations of prefrontal D2R and SERT with the analyzed RSNs indicate that the interplay of DA and 5-HT is likely to be paramount to medial prefrontal function and to the RSNs modulated by it, primarily the DMN.

Future studies will be required to further investigate the mechanisms underlying the observed correlations. Importantly, as opposed to neurotransmitters such as the mainly excitatory glutamate or the mainly inhibitory GABA, DA and 5-HT modulate brain function heterogeneously [55]. The heterogeneity of the modulatory effects may stem from the various DA and 5-HT receptor subtypes interacting with glutamate and GABA in differing manners and their different distributions across the brain. For example, in the case of 5-HT, presynaptic 5-HT_1A_, 5-HT_1B_ and 5-HT_6_ receptors have been shown to decrease glutamate release, while 5-HT_3_ receptors increase the release of glutamate. 5-HT_2_ receptors increase GABA release and can reduce or enhance glutamate release depending on the region [55]. Intriguingly, several studies have suggested a relationship between DA and 5-HT in disease, reward and addiction [56, 57]. Anatomically, it has been shown that the raphé nuclei send serotonergic projections to the ventral tegmental area [57, 58], while in turn receiving top-down afferents from the prefrontal cortex [53] and other areas [59, 60]. Additionally, the VTA also receives top-down input from the mPFC [60, 61]. In our study, additional analysis showed medial prefrontal SERT availability correlated positively with the short-path rs-FC between VTA and MB. This finding hints towards the role of serotonergic prefrontal top-down modulation on the relationship between the raphé nuclei and the VTA and conversely may represent one of the ways of interaction between serotonergic and dopaminergic neurotransmission. Such complex loops mediated by several regions and neurotransmitters may be further elucidated by PET/fMRI studies employing several tracers in the same cohort.

### Limitations and general remarks

Our study is the first exploring the correlation of molecular receptor and transporter availability with RSNs using a simultaneous PET/fMRI approach in rats. As the data were acquired under anesthesia, this effect needs to be taken into account for the interpretation of results. While the anesthesia was kept at levels recommended previously [62] and shown to enable stable physiological readouts [63], some confounding effects cannot be excluded for either of the fMRI [62], [^11^C]raclopride [64] or [^11^C]DASB readouts [65]. However, performing such experiments in small laboratory animals opens up the great opportunity to study such interactions under very controlled conditions and maximized cohort uniformity. Factors such as nutrition, lifestyle, age or gender known to impact D2R and SERT availabilities in a regionally specific manner [66, 67] can be excluded when interpreting the observed correlations, thereby enabling an inherently complementary readout to human studies.

Furthermore, some of the correlations presented in our study are moderate and did not survive FDR correction. Two factors may represent possible causes for this issue. First, compared to large clinical studies, the sizes of our cohorts are relatively limited. Second, D2R and SERT densities are not the sole modulators of rs-FC, other neurotransmitters and receptor types probably having as of yet undiscovered associations with rs-FC. Therefore, our study sheds light on a part of the picture of interactions between neurotransmitter systems and rs-FC; similarly designed studies are still required to thoroughly elucidate this aspect. Importantly, PET/fMRI offers the possibility to generate multi-level data on this very complex matter. In future, a PET/fMRI database, similar to already existing fMRI databases, may be of interest for potential large cohort meta-analyses to this extent. Since most psychiatric medications aim to normalize brain function by interacting with certain receptors or transporters, applying novel analysis methods, as well as machine learning approaches to this type of data can help understand the link between molecular changes and functional changes in the brain, enable the accurate prediction of drug therapies and improve development of treatment strategies for psychiatric disorders.

## Conclusion

Here we present that the local availability of D2R and SERT have regionally specific fingerprints on RSNs. We apply a novel analysis method of simultaneously acquired PET/fMRI data in rats which enables to investigate the modulatory role of neurotransmitter systems on rs-FC at baseline levels. Further studies exploring the correlations of other neurotransmitter systems such as norepinephrine with rs-FC will be of great value to elucidate their respective influence on brain function. Future similarly designed studies may improve the general understanding of brain function on several levels, as well as the development of novel drug therapies for several psychiatric diseases.

## Materials and Methods

### Animals

Male Lewis rats (n = 59) provided by Charles River Laboratories (Sulzfeld, Germany) were divided into two cohorts for [^11^C]raclopride (365 ± 49 g, n = 29) and [^11^C]DASB (354 ± 37 g, n = 30, see *Supplementary Figure 1* for the rat weights). These weights corresponded to ages of approximately 15 weeks. The rats were kept on a 12-hour day-night cycle at a room temperature of 22 °C and 40-60% humidity and received standard chow food and water *ad-libitum*. All experiments were conducted according to the German federal regulations regarding use and care of experimental animals and were approved by the local authorities (Regierungspräsidium Tübingen).

Initially, a total of 50 rats were scanned using [^11^C]DASB and 37 rats were scanned using [^11^C]raclopride. 20 scans of the [^11^C]DASB cohort and 8 scans of the [^11^C]raclopride cohort had to be excluded due to either motion, insufficient tracer specific activity, paravenous catheters or image artifacts.

### Radiotracer synthesis

For a detailed account of radiotracer synthesis, please refer to *Supplementary Methods*.

The radioactive tracers had molar activities of 83 ± 29 GBq/µmol for [^11^C]raclopride 57 ± 37 GBq/µmol for [^11^C]DASB at the start of the PET acquisition (*Supplementary Figure 2B and C*).

### Simultaneous PET/MRI experiments

Anesthesia was induced in knock-out boxes by delivering 3 % isoflurane in regular air until reflex tests indicated sufficient sedation. For the following preparation steps the concentration of isoflurane was reduced to 2 %. The weights of the animals were measured and a catheter was placed into a tail vein using a 30 G needle for tracer administration. Subsequently, the rats were transferred onto a dedicated feedback temperature-controlled rat bed (Medres, Cologne, Germany). A rectal probe was positioned to monitor and maintain a stable body temperature at 36.5° C and a breathing pad was used to observe respiration rates. Finally, the animals were introduced into the PET/MRI scanner and the isoflurane concentration was reduced to 1.3 % during the scan.

The scans were acquired using a small-animal 7 T ClinScan scanner (Bruker BioSpin MRI, Bruker, Ettlingen, Germany) with a linearly polarized RF coil (Bruker) of 72 cm in diameter for transmission and a four channel rat brain coil (Bruker) for reception. Localizer scans were first acquired to accurately position the rat brains into the center of the PET/MRI field of view. Subsequently, local field homogeneity was optimized by measuring local magnetic field maps. Anatomical reference scans were then performed using T2-weighted MRI sequences (TR: 1800 ms, TE: 67.11 ms, FOV: 40 x 32 x 32 mm^3^, image size: 160 x 128 x 128 px, Rare factor: 28, averages: 1). Finally, T2*-weighted gradient echo EPI sequences (TE: 18 ms, TR: 2500 ms, 0.25 mm isotropic resolution, FoV 25 x 23 mm^2^, image size: 92 x 85 x 20 px, slice thickness: 0.8 mm, 20 slices) were acquired for functional MR imaging.

A small-animal PET insert developed in cooperation with Bruker (Bruker Biospin, Ettlingen Germany) was used for [^11^C]DASB and [^11^C]raclopride acquisitions. This insert is the second generation of a PET insert developed in-house described previously [68]. Both PET inserts have similar technical specifications. The radioactive tracers were applied via a bolus plus constant infusion protocol with a K_bol_ of 38.7 minutes using an initial bolus of 341 ± 65.2 MBq for [^11^C]raclopride and 152 ± 44 MBq for [^11^C]DASB in a volume of 0.48 ml over 20 seconds, followed by a constant infusion of 15 µl/min until the end of the scan. PET/fMRI acquisition was started simultaneously with the tracer injection and was performed over a period of 80 minutes. The PET data were saved as list-mode files and reconstructed using an ordered-subsets expectation maximization 2D (OSEM-2D) algorithm written in-house.

### Data preprocessing

Preprocessing was performed using Statistical Parametric Mapping 12 (SPM 12, Wellcome Trust Centre for Neuroimaging, University College London, London, United Kingdom) and Analysis of Functional NeuroImages (AFNI, National Institute of Mental Health (NIMH), Bethesda, Maryland, USA) as reported previously [69]. In addition, we added a nuisance removal based on a method reported elsewhere [70]. First, all fMRI scans were realigned using SPM and the three translation and three rotation motion parameters were stored. Additionally, mean images were created for all scans and used to create binary masks using AFNI. Additional binary brain masks were created for the T2-weighted anatomical MRI reference scans and the reconstructed PET scans. The masks were applied for brain extraction from all mentioned datasets. For fMRI, images containing extra-cerebral tissue were also created for later use. The skull-stripped PET and fMRI scans were then coregistered to their respective anatomical references. Afterward, the anatomical reference scans were used to calculate spatial normalization parameters to the Schiffer rat brain atlas and the obtained normalization parameters were applied to the fMRI and PET datasets. Coregistration and normalization were visually evaluated for each subject and every modality. Then, nuisance removal was performed for the fMRI scans. To this extent, a multiple linear regression model was applied containing the six motion parameters stored after initial realignment, as well as the first 10 principal components of the signal extracted from the images containing extra-cerebral tissues, as described by Chuang et al. [70]. Finally, a 1.5 x 1.5 x 1.5 mm^3^ full-width-half-maximum Gaussian kernel was applied to all fMRI and PET datasets for spatial smoothing [71].

### Data analysis

In the following, an overview of the analysis of the preprocessed data is provided. A graphical description of the analysis pipeline used is shown in Figure 1.

### Functional MRI data analysis

Resting-state functional connectivity was calculated using a seed-based approach in the interval from 40 to 80 minutes after scan start to ensure tracer equilibrium for PET (please refer to PET data analysis section). To this extent, 28 regions comprising the DMN, SN and SMN were selected from the Schiffer rat brain atlas (a list of the regions is provided in *Supplementary Table 1*). The SPM toolbox Marseille Boíte À Région d’Intérêt (MarsBaR) was employed to extract fMRI time-courses from all regions [72]. These were then used to calculate pairwise Pearson’s r correlation coefficients for each dataset, generating correlation matrices containing 28 x 28 elements. Self-correlations were set to zero. The computed Pearson’s r coefficients then underwent Fischer’s transformation into z values for further analysis.

Several rs-FC metrics were computed to quantify the properties of the analyzed networks. In addition to edge-wise rs-FC, regional node strengths were calculated as the sum of all correlations of one node to the regions belonging to the same network. Inter-network node strengths were defined as the sum of the correlations of one node to the regions of another network. On a network level, within-network strengths were defined as the sum of all edges comprising a network. Between-network strengths were calculated as the sum of all correlations between two sets of regions belonging to two networks [73].

For a detailed report of these steps please refer to *Supplementary Methods*.

### PET data analysis

Static PET scans were reconstructed from 40 to 80 minutes after the start of the PET data acquisition to ensure tracer equilibrium between target and reference region. To enhance signal-to-noise ratios and due to the negligible differences in tracer uptake between left and right hemispheres, each bilateral region of the Schiffer rat brain atlas was merged to one volume of interest (VOI). Following preprocessing, tracer uptake values of the 27 generated VOIs were calculated for each dataset. Binding potentials (BP_ND_) values were computed from the DVR-1 (equation 1) using the whole cerebellum as a reference region for [^11^C]raclopride and the cerebellar grey matter as a reference region for [^11^C]DASB [74, 75].

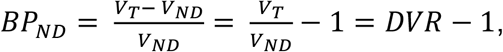

where:

- *BP*_*ND*_ is the binding potential
- *V*_*T*_ is the total volume of distribution
- *V*_*ND*_ is the volume of distribution in a reference tissue
- *DVR* is the distribution volume ratio

For the generation of BP_ND_ maps, the above equation was applied for each voxel, where *V*_*ND*_ was defined as the mean uptake of all voxels included in the reference region, while *V*_*T*_ was the uptake of each respective voxel, resulting in single BP_ND_ value for each voxel in every subject. Using the subject BP_ND_ maps group-level BP_ND_ maps were calculated for both cohorts. For correlation analyses, VOI-based BP_ND_ values were calculated, *V*_*T*_ representing the mean uptake of all voxels comprised by the respective VOI. Previous similar studies reported adjusting BP_ND_ values for mean global signal to control for global effects [5]. The mentioned study indicated the high inter-individual correlations between BP_ND_ values of different regions. Here, we reproduced the finding by calculating the correlations between regional BP_ND_ values and confirmed this observation (please refer to *Supplementary Information* for an exemplary correlation analysis between [^11^C]DASB bindings in the mPFC and CPu). Thus, since our aim was to elucidate the correlations of rs-FC with the distributions of either D2R or SERT binding between different regions, in our study individual BP_ND_ values also underwent a global normalization for each dataset to discard such effects, generating normalized BP_ND_ values (BP_ND-norm_), as described in the study mentioned above [5].

### PET/fMRI data analysis

To investigate the influence of [^11^C]raclopride and [^11^C]DASB in the CPu and mPFC on rs-FC, we evaluated their relationships between BP_ND-norm_ values and rs-FC measures described above. Inter-individual correlations between regional BP_ND-norm_ values and rs-FC metrics were calculated using Pearson’s r. This procedure was performed between each regional BP_ND-norm_ and every rs-FC metric described above to determine potential correlations between edges, regional node strengths, inter-regional node strengths, within-network strengths and between-network strengths and regional D2R or SERT densities. Additionally, the computed correlations between BP_ND-norm_ and each rs-FC metric were tested for statistical significance and a false discovery rate (FDR) correction was performed for a threshold of 0.05 using the Benjamini-Hochberg procedure.

## Acknowledgements

We thank Dr. Julia Mannheim, Dr. Rebecca Rock, Dr. Neele Hübner, Dr. Andreas Dieterich, Ines Herbon, Stacy Huang, Funda Cay, Linda Schramm and Sandro Aidone (Werner Siemens Imaging Center, Department of Preclinical Imaging, University of Tuebingen) for their technical and adminstrative support. We thank the Radiopharmacy department (Werner Siemens Imaging Center, University of Tuebingen) for the tracer production. This study is also part of the PhD thesis of Tudor Ionescu.

## Author contributions

- Conceptualization: TI, BBB, HFW
- Methodology: TI, RH
- Software: TI, RH
- Validation: TI
- Formal analysis: TI
- Investigation: TI, MA
- Resources: BJP
- Data curation: TI
- Writing – original draft: TI
- Writing – review and editing: MA, RH, BBB, AM, BJP, HFW, KH
- Visualization: TI, KH
- Supervision: BJP, HFW, BBB, KH
- Project administration: BJP, HFW, BBB, KH
- Funding acquisition: BJP, HFW, BBB

## Data availability

The original dataset will be made available upon request.

