## supplemental information for "Striatal and prefrontal D2R and SERT distributions contrastingly correlate with default-mode connectivity"

### Supplementary Methods

#### Schiffer rat brain atlas

Brain parcellation was performed based on the Schiffer rat brain atlas. A total of 14 regions were selected. In **Supplementary Table 1** a list of the regions, their volumes and abbreviations is provided.

**Supplementary Table 1: Regions selected for analysis from the Schiffer brain atlas, along with their volumes and abbreviations.**

| Brain region (ROI) | ROI volume [mm$^3$] | Abbreviation |
|---|---|---|
| Nucleus Accumbens | 7.944 | NAc |
| Amygdala | 21.120 | Amyg |
| Caudate Putamen | 43.552 | CPu |
| Cingulate Cortex | 14.480 | Cg |
| Insular Cortex | 21.128 | Ins |
| Medial Prefrontal Cortex | 6.304 | mPFC |
| Motor Cortex | 32.608 | MC |
| Orbitofrontal Cortex | 18.936 | OFC |
| Parietal Cortex | 7.632 | PaC |
| Retrosplenial Cortex | 18.920 | RS |
| Somatosensory Cortex | 71.600 | S1 |
| Anterodorsal Hippocampus | 25.064 | CA1 |
| Posterior Hippocampus | 9.784 | CA1-p |
| Thalamus | 30.712 | Th |

**Supplementary Table 2** lists the regions comprising each of the analyzed resting-state networks.

**Supplementary Table 2: Regions comprising every analyzed resting-state network.**

| Default-mode network (DMN) | Salience network | Sensory-motor network |
|---|---|---|
| mPFC | NAc | CPu |
| OFC | Amyg | M1 |
| PaC | Cg | S1 |
| RS | Ins | Th |
| CA1 | | |
| CA1-p | | |

**Radiotracer synthesis**

[$^{11}$C]Raclopride synthesis was performed as described by Langer et al. [76] with the optimized HPLC conditions of van Laeken et al. [77]. After dilution with water, the isolated fraction was trapped on a conditioned Strata-X cartridge (Phenomenex, Aschaffenburg, Germany), eluted with 0.5 ml ethanol and diluted with 5 ml phosphate-buffered saline. [$^{11}$C]DASB was synthesized similar to the procedure reported by Wilson et al. [78]. Briefly, [$^{11}$C]MeI was trapped in a solution of 2 mg precursor in 500 µl DMSO. After heating to 100 °C for 2 min the reaction was diluted with 1.5 ml HPLC eluent (3 mM $Na_2HPO_4$ containing 64 % MeCN) and purified on a Luna C18(2) column (250 mm x 10 mm, Phenomenex). The isolated peak was diluted with 70 ml water containing 20 mg sodium ascorbate, loaded onto a conditioned Strata-X cartridge (Phenomenex), eluted with 0.5 ml ethanol and diluted with 5 ml phosphate-buffered saline.

The radioactive tracers had molar activities of 82 ± 28 GBq/µmol for [$^{11}$C]raclopride 50 ± 16 GBq/µmol for [$^{11}$C]DASB at the start of the PET acquisition (Supplementary Figure 2B and C).

**Seed-based resting-state functional connectivity analysis**

To derive rs-FC between the selected regions the regional time-series of the preprocessed fMRI scans were extracted using the MarsBaR toolbox. Every regional BOLD time-course was computed as the mean of all the time-courses of the voxels comprising the respective region:

$$Y_I = \frac{1}{N_I} \sum_{i \epsilon I} y_i \quad , \tag{1}$$

where $I$ is the delineated brain ROI, $Y_I$ its derived time-series, $N_I$ is the number of voxels comprising ROI $I$ and $y_i$ is the time series of voxel $i$.

Pearson's r correlation coefficients were calculated between each the extracted signals from each pair of regions to determine seed-based rs-FC employing an algorithm written in-house:

$$r = \frac{\sum_{i=1}^{n}(u_i - \bar{u})(v_i - \bar{v})}{\sqrt{\sum_{i=1}^{n}(u_i - \bar{u})^2}\sqrt{\sum_{i=1}^{n}(v_i - \bar{v})^2}} \quad , \tag{2}$$

where $r$ is Pearson's correlation coefficient, $n$ is the number of samples, $u_i$ and $v_i$ are the single samples and $\bar{u}$ and $\bar{v}$ are the arithmetical sample means.

Since the cerebellum was excluded from fMRI analysis, as it was used as reference region for the determination of $BP_{nd}$ values, the approach yielded individual 48 x 48 correlation matrices. The 48 self-correlations were set to zero; thus, each individual scan included 2256 pair-wise correlation coefficients.

For group-level analyses, the calculated correlation coefficients were first transformed into Fischer's z-scores:

$$Z = \frac{1}{2} \ln \frac{(1+r)}{(1-r)} = \mathrm{arctanh}(r) \quad , \tag{3}$$

Each z-score was averaged over the two cohorts respectively and then retransformed to the interval [−1, 1] to be used for further analyses.

#### Supplementary Results

The bodyweights of the scanned animals and the radioactivity of the injected tracers are depicted in **Supplementary Figure 1**.

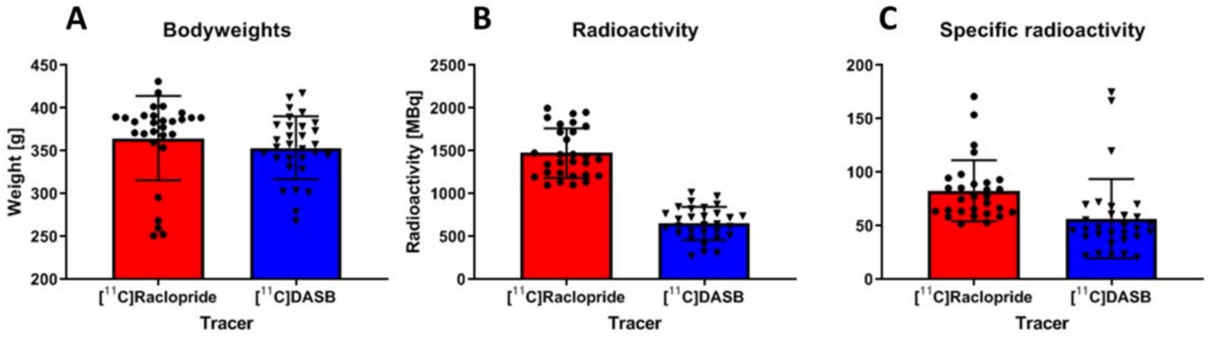


**Supplementary Figure 1: (A) Bodyweights of the used rats, (B) radioactivity and (C) specific radioactivity values of injected tracers (n = 29 for [$^{11}$C]raclopride and n = 30 for [$^{11}$C]DASB).**

The distributions of the injected tracers, as well as the three RSNs investigated are presented in **Supplementary Figure 2**.

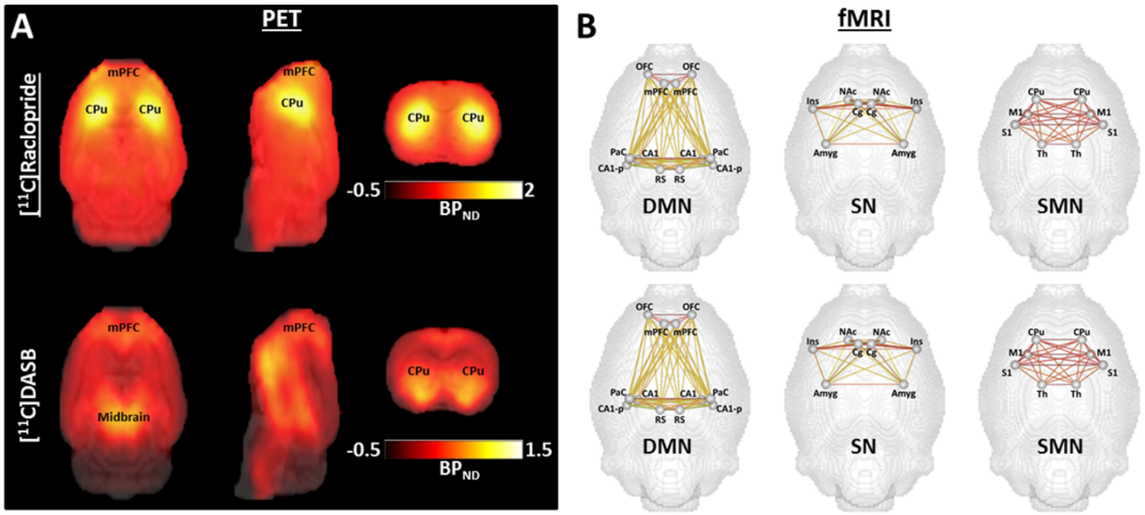

.

**Supplementary Figure 2: PET and fMRI analyses. (A)** Group-mean [$^{11}$C]raclopride and [$^{11}$C]DASB $BP_{ND}$ maps. **(B)** Resting-state networks extracted from the mean rs-FC analysis performed for the [$^{11}$C]raclopride (upper row) and [$^{11}$C]DASB (lower row) cohorts. Connectivity patterns are presented for the default-mode network, salience network and sensory-motor network. Abbreviations: DMN = default-mode network; SN = salience network; SMN = sensorimotor network. Please refer to *Supplementary Table 2* for a list of regions comprising each network.

An exemplary scatter plot between the [$^{11}$C]DASB $BP_{ND}$ values of the CPu and mPFC is shown in **Supplementary Figure 3**.

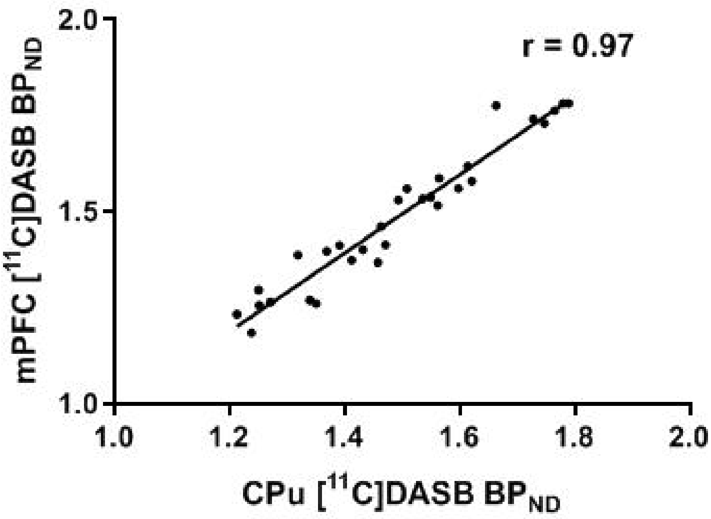


**Supplementary Figure 3: Scatter plot between the [$^{11}$C]DASB $BP_{ND}$ values in the mPFC and CPu.**

The scatter plots corresponding to the correlations between all networks presented in the main manuscript are depicted in **Supplementary Figure 4** - **Supplementary Figure 7**.

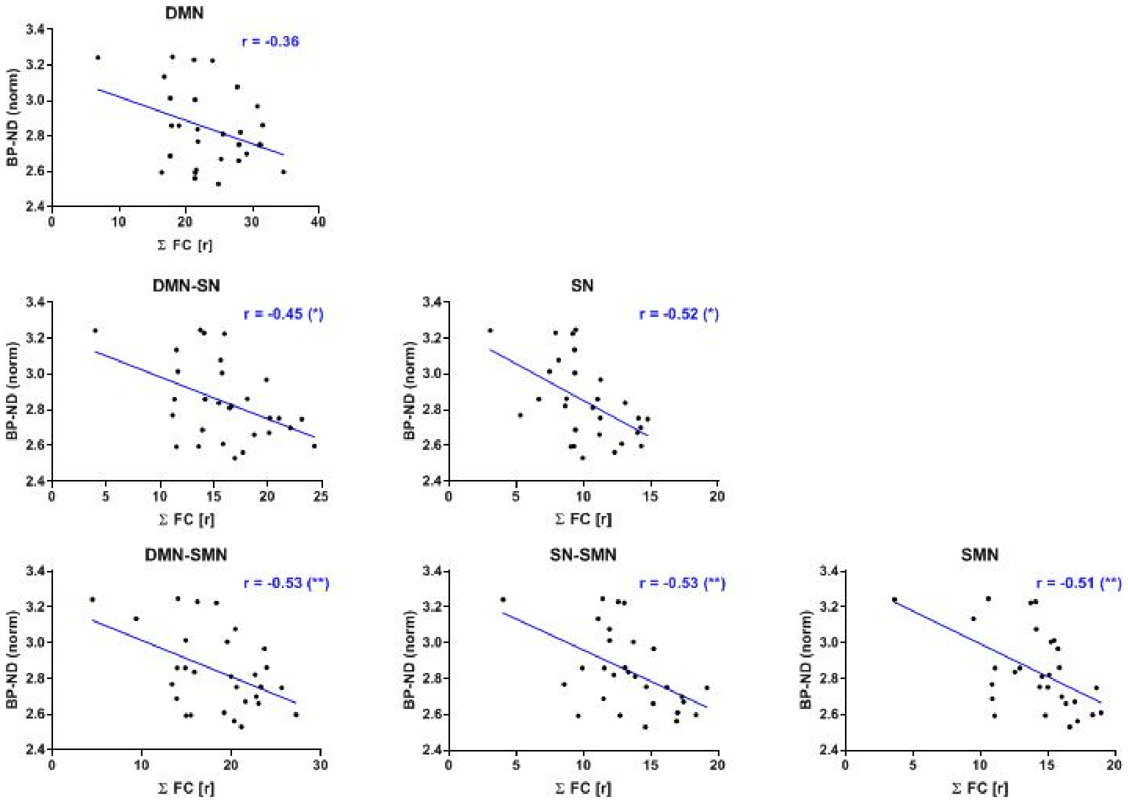


**Supplementary Figure 4: Scatter plots depicting the correlations between CPu [[11]C]raclopride $BP_{ND\text{-}norm}$ values and within and between-network rs-FC.** Abbreviations: DMN = default-mode network, SN = salience network, SMN = sensorimotor network.

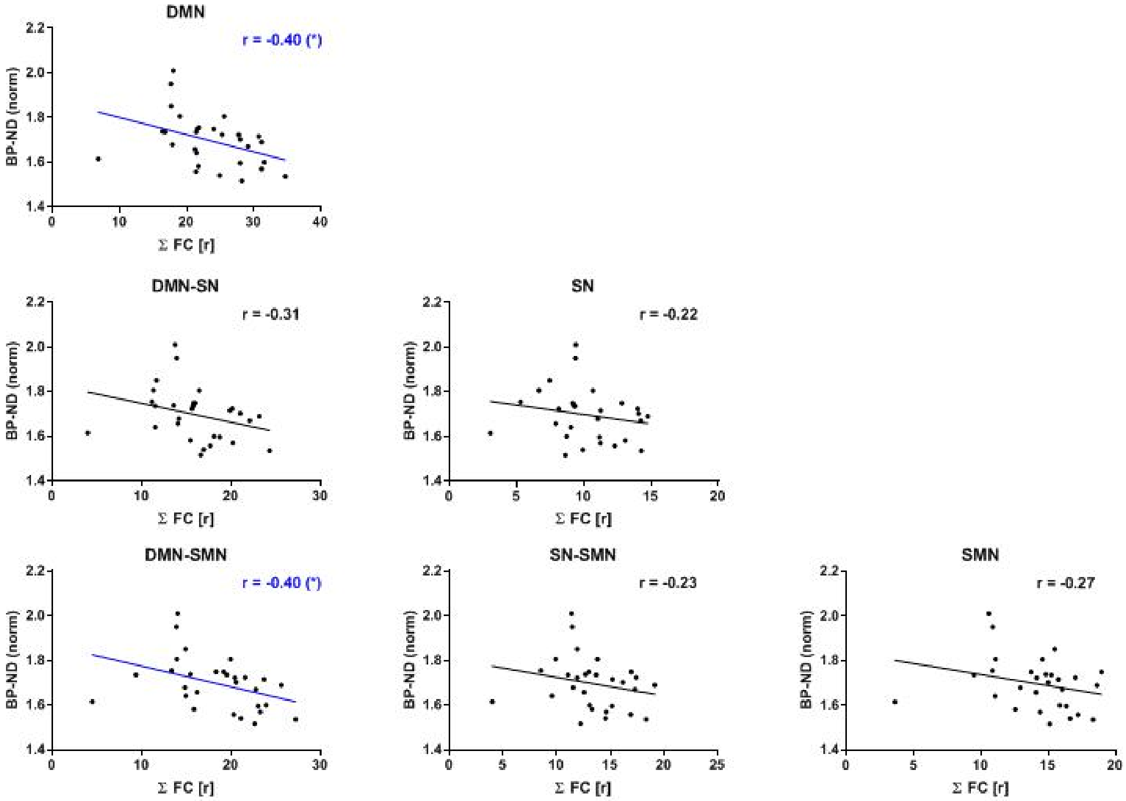


**Supplementary Figure 5: Scatter plots depicting the correlations between mPFC $[^{11}C]$raclopride $BP_{ND\text{-}norm}$ values and within and between-network rs-FC.** Abbreviations: DMN = default-mode network, SN = salience network, SMN = sensorimotor network.

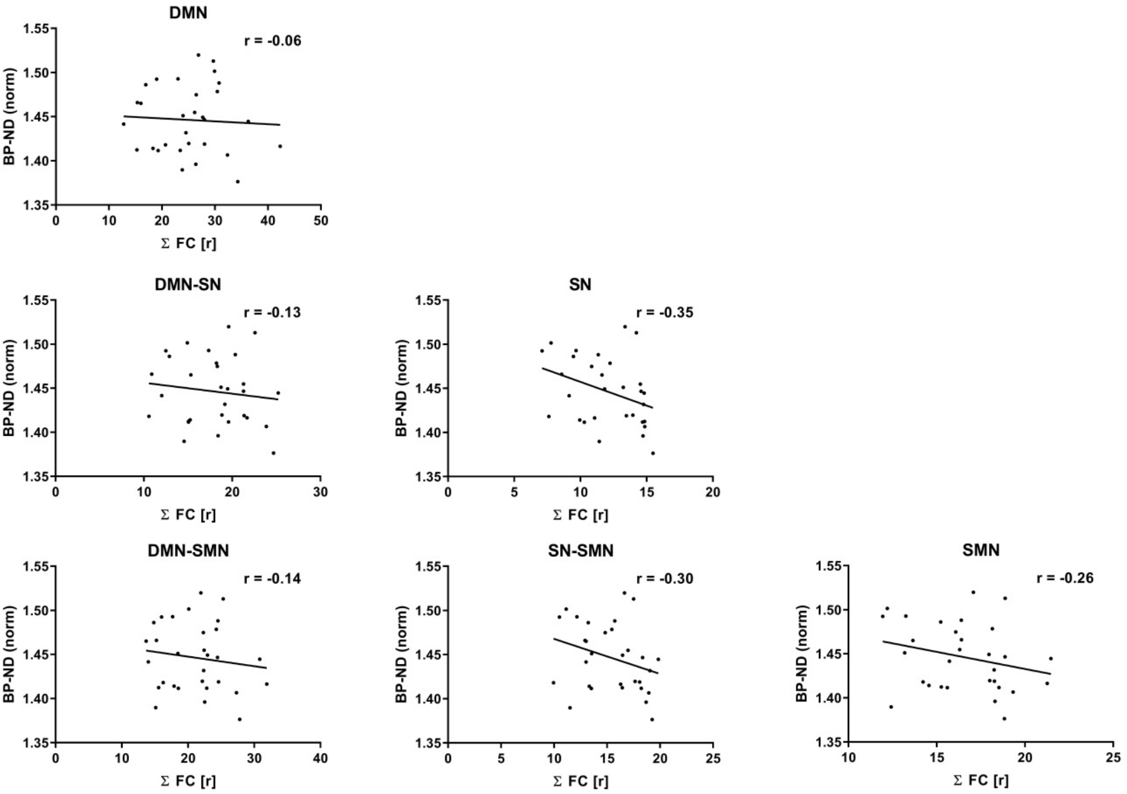


**Supplementary Figure 6: Scatter plots depicting the correlations between mPFC $[^{11}C]$DASB $BP_{ND-norm}$ values and within and between-network rs-FC.** Abbreviations: DMN = default-mode network, SN = salience network, SMN = sensorimotor network.

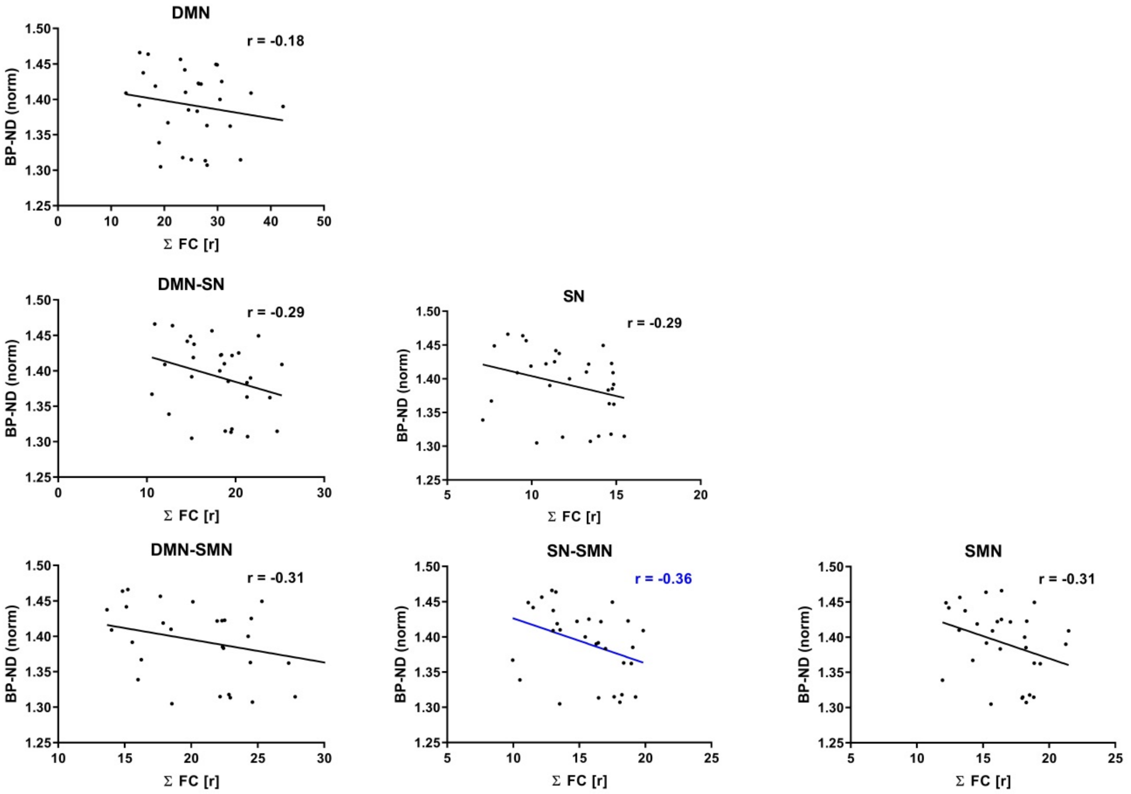


**Supplementary Figure 7: Scatter plots depicting the correlations between mPFC $[^{11}C]$DASB $BP_{ND\text{-}norm}$ values and within and between-network rs-FC.** Abbreviations: DMN = default-mode network, SN = salience network, SMN = sensorimotor network.

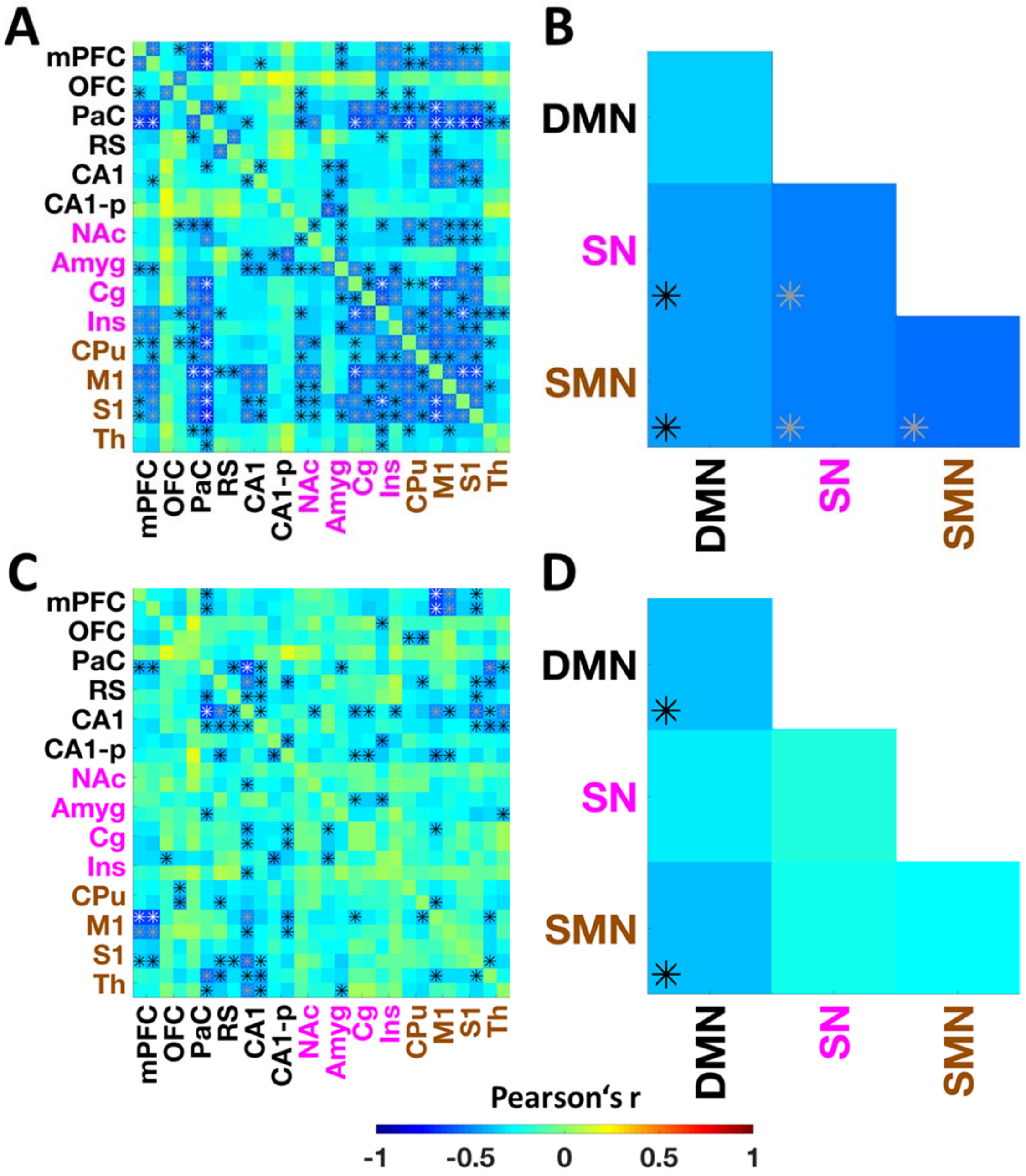


**Supplementary Figure 8: Significant correlations for CPu (A and B) and mPFC (C and D) [$^{11}$C]raclopride bindings and rs-FC shown at uncorrected thresholds of $p < 0.05$ (black asterisks), $p < 0.01$ (grey asterisks) and $p < 0.001$ (white asterisks).**

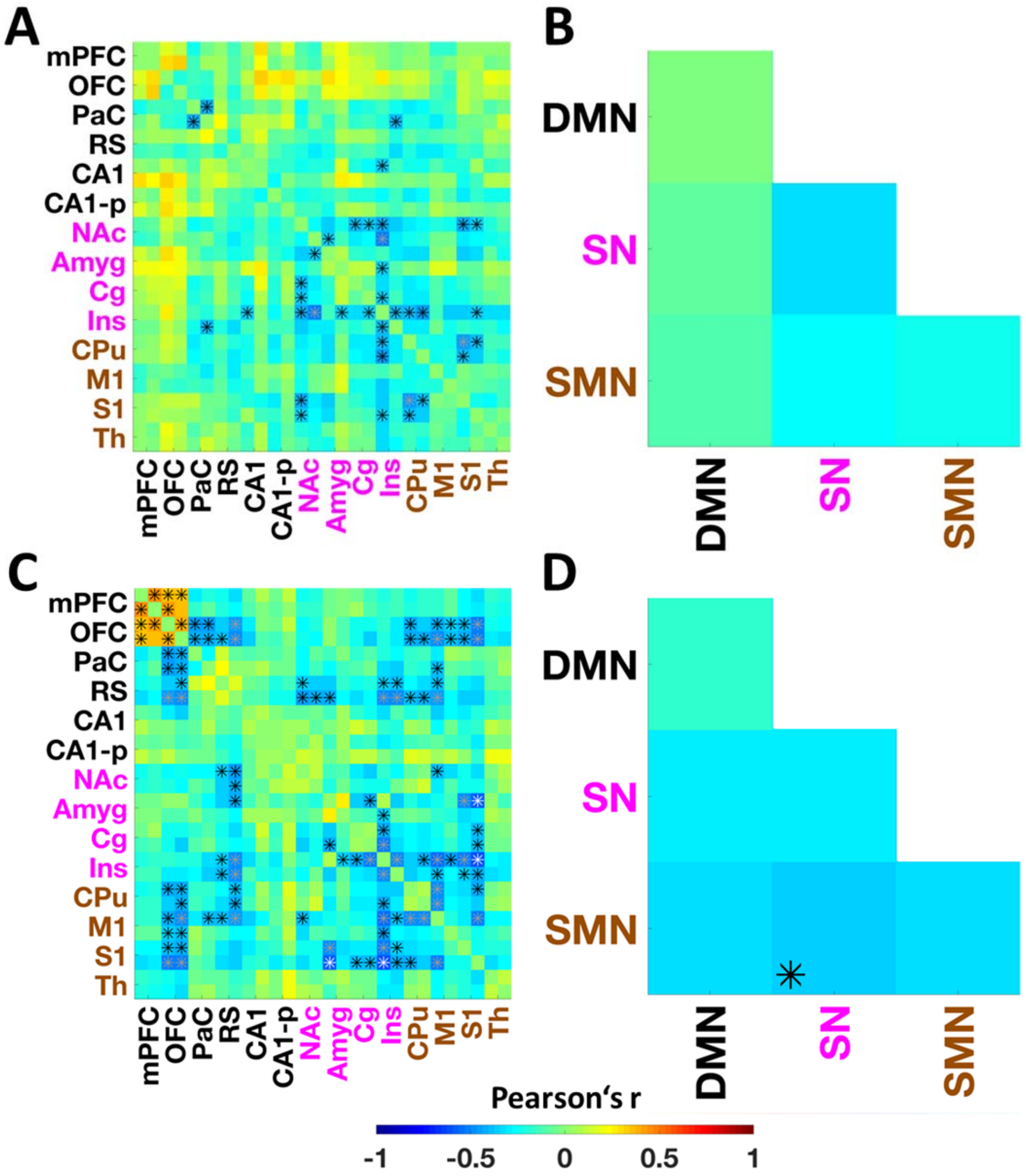


**Supplementary Figure 9: Significant correlations for CPu (A and B) and mPFC (C and D) $[^{11}C]$DASB bindings and rs-FC shown at uncorrected thresholds of $p < 0.05$ (black asterisks), $p < 0.01$ (grey asterisks) and $p < 0.001$ (white asterisks).**

We performed an additional analysis to further emphasize the effect according to which mPFC SERT binding positively correlates with rs-FC within the anterior DMN in contrast to rs-FC between the anterior DMN and posterior DMN, SN and SMN, shifting the balance to anterior DMN processing (**Supplementary Figure 10**).

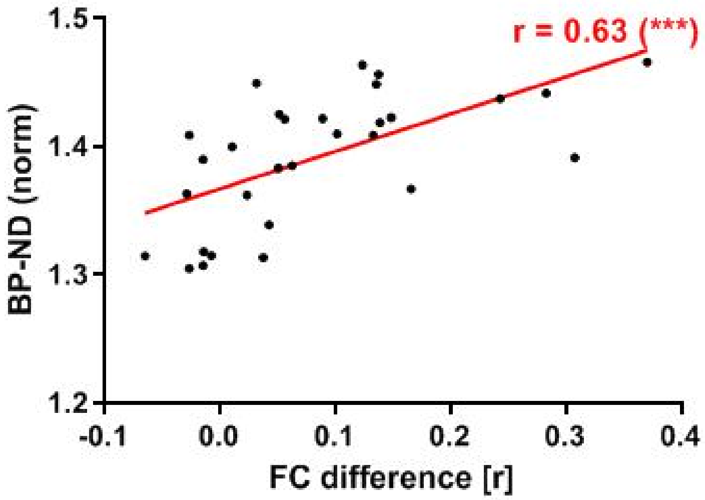


**Supplementary Figure 10: Correlation between [$^{11}$C]DASB binding in the mPFC and the difference of the average connectivity within the anterior DMN, comprising all edges between mPFC and OFC, and the rs-FC between the anterior DMN and the posterior DMN, SN and SMN was highly significant (r = 0.63, p = 0.0003).**
